# Near-surface wind variability over spatiotemporal scales relevant to plume tracking insects

**DOI:** 10.1101/2023.01.18.524580

**Authors:** Jaleesa Houle, Floris van Breugel

**Affiliations:** Department of Mechanical Engineering, University of Nevada, Reno

**Keywords:** near surface wind, odor plume tracking, microscale wind dynamics

## Abstract

Odor plume tracking is an important biological process for many organisms, and flying insects have served as popular model systems for studying these behaviors both in the field and in lab settings. The shape and statistics of the airborne odor plumes that insects follow are largely governed by the wind that advects them. Prior atmospheric studies have investigated aspects of microscale wind patterns with an emphasis on characterizing pollution dispersion, enhancing weather prediction models, and for assessing wind energy potential. Here, we aim to characterize microscale wind dynamics through the lens of short-term ecological functions by focusing on spatial and temporal scales most relevant to an insect actively searching for an odor source. We collected and compared near-surface wind data across three distinct environments (sage steppe, forest, and urban) in locations across Northern Nevada. Our findings show that near-surface wind direction variability decreases with increasing wind speeds and increases in environments with greater surface complexity. Across environments, there is a strong correlation between the variability in wind speed (i.e. turbulence intensity) and wind direction (i.e. the standard deviation in wind direction). In some environments, the standard deviation in wind direction varied as much as 15° to 75° on time scales of 1-10 minutes. We draw insights between our findings and previous plume tracking experiments to provide a general intuition for future field research and guidance for wind tunnel experimental design. From our analyses, we hypothesize that there may be an ideal range of wind speeds and environment complexity in which insects will be most successful when tracking an odor plume to its source.

## 1. Introduction

Across many species and ecosystems, the ability to successfully localize viable food sources or mates through olfactory search is critical for their survival. From shrimp to sharks [1, 2] and moths to mammals [3, 4, 5], there are countless examples of aquatic, terrestrial, and aerial animals that rely on the synergy of multiple sensory modalities to track odor plumes to their sources. The sensory experience of a plume tracking animal is determined by their movement as well as the shape and statistics of the plume, which are in turn governed by characteristics of the fluid medium responsible for its advection. In laminar flow, odor plumes take on a simple ribbon-like shape [6, 3], whereas plumes in more turbulent flow become more dispersed and broken up [7, 8, 9]. Field studies in turbulent air flow with moths [10] and fruit flies [11] suggest that insects can still be quite successful plume trackers on spatial scales of 10-100 meters and time scales of tens of minutes despite the flow complexity caused by turbulence. To better understand both flying insects’ plume tracking algorithms and the ecological pressures that have driven their evolution, it is important that we characterize the spatial and temporal statistics of natural wind on the scale of tens of minutes across 10-100 meters, and in different environments that flying insects encounter.

The study of wind is quite broad, with applications spanning from the lowest portion of the atmosphere (surface roughness sublayer) through to higher altitudes. First introduced in 1954, Monin-Obukhov Similiarity Theory (MOST) is widely used as a general framework for quantifying atmospheric stability and turbulence [12]. Although MOST is often used to describe the vertical wind profile spanning up to the lower layer of the troposphere, this relationship does not hold within the surface roughness sublayer, posing challenges for successfully classifying near-surface wind conditions in highly vegetated or urban areas. [13, 14]. The challenge of classifying near-surface wind conditions is often compounded in microscale environments (*<* 1 square kilometer) due to increasing surface heterogeneity caused by land development [15, 16]. Efforts have been made toward developing a mathematical framework for describing the wind profile within plant canopies [17, 18], however, these frameworks typically assume a homogeneous environment and require measurement of heat flux induced by vegetation. Furthermore, plant canopy wind models are not generally applicable in other environments such as cities or sparse-vegetation landscapes.

Prior atmospheric studies have sought to quantify near-surface wind patterns for applications in weather modeling, pollution dispersion, and wind energy production. Near-surface nocturnal wind regimes have been of particular interest due to the higher occurrences of submeso motions and other flow phenomena, which become more readily observable in strongly stable nocturnal atmospheric conditions [19, 20, 21]. [22] and [23] assess data collected from multiple environments and describe temporal changes of wind direction and speed variability over scales of approximately 10-15 minutes in nocturnal conditions. These studies found that standard deviations in wind direction are inversely related to wind speed with directional variability highest in wind speeds less than 2 m/s. There are undoubtedly many nocturnal insect species for which these prior studies would be applicable.

Near-surface wind regimes in complex urban areas are also of particular interest to researchers seeking to improve existing pollution models [24, 25, 26]. Both [25] and [26] used an array of wind sensors distributed across various heights and distances within areas of less than 1km to assess patterns in urban wind variability. Since wind profiles in the urban environment are not well understood, these works explored statistical quantities at and above street level. [25] found horizontal and vertical turbulence intensities measurements from two different cities to be in general agreement, suggesting that near-surface turbulent characteristics may be similar across urban areas. Other case studies have examined wind profiles specific to an area of interest for wind energy assessment [27, 28, 29]. Though these efforts have all contributed to our collective knowledge of near-surface wind patterns, we still do not have a comprehensive understanding of the fine-scale wind direction and wind speed variability which occurs across environments of varying surface complexity in daytime convective processes. Here, we seek to compare short-term wind variability across three distinct environments as a means of gaining intuition on the key differences an insect might experience while actively tracking an odor plume.

To gain insight into daytime microscale wind conditions, we collected near-surface wind data in three qualitatively distinct environments in Northern Nevada (playa/sage steppe, urban, and forest) and statistically quantified temporal and spatial variability in direction and speed. Since our goal was to understand the significance of wind variability over scales relevant to a plume tracking insect, we focused our analysis on wind patterns occurring over periods of less than 15 minutes, across distances of less than 200 meters, and at an altitude of 2 meters above the ground. Prior field research has documented male gypsy moths flying within 1 meter from the ground while tracking female pheromones [30], and other works suggest that insects such as *Drosophila* and honeybees adjust their flight altitude based on wind speed in order to maintain constant optic flow [31]. Though the average altitude that insects inhabit may vary based on environment, comparing wind observations at a height of 2 meters above ground level across environments provides insight on the sensory information that a wide variety of insects may be likely to experience while actively tracking an odor. Our findings indicate that near-surface wind patterns over these spaces and time ranges are most consistently variable in low wind speeds and in environments with higher surface complexity.

## 2. Methods

### 2.1 Site Descriptions

Wind data was recorded across various locations in Northern Nevada (Figure 1). Three qualitatively distinct environments were chosen for this work: playa/sage steppe, forest, and urban areas. The playa environment (Black Rock Desert) can be characterized as open, flat, and sandy with no vegetation. The sage steppe environment (Lemmon Valley) is an open area with surrounding hills approximately 0.5 to 1 kilometer away, with an average vegetation height of approximately 0.75 meters. Qualitatively, we group these environments together as “open” landscapes, though we note the distinction in wind profiles due to differences in surface roughness elements.

**Figure 1:**
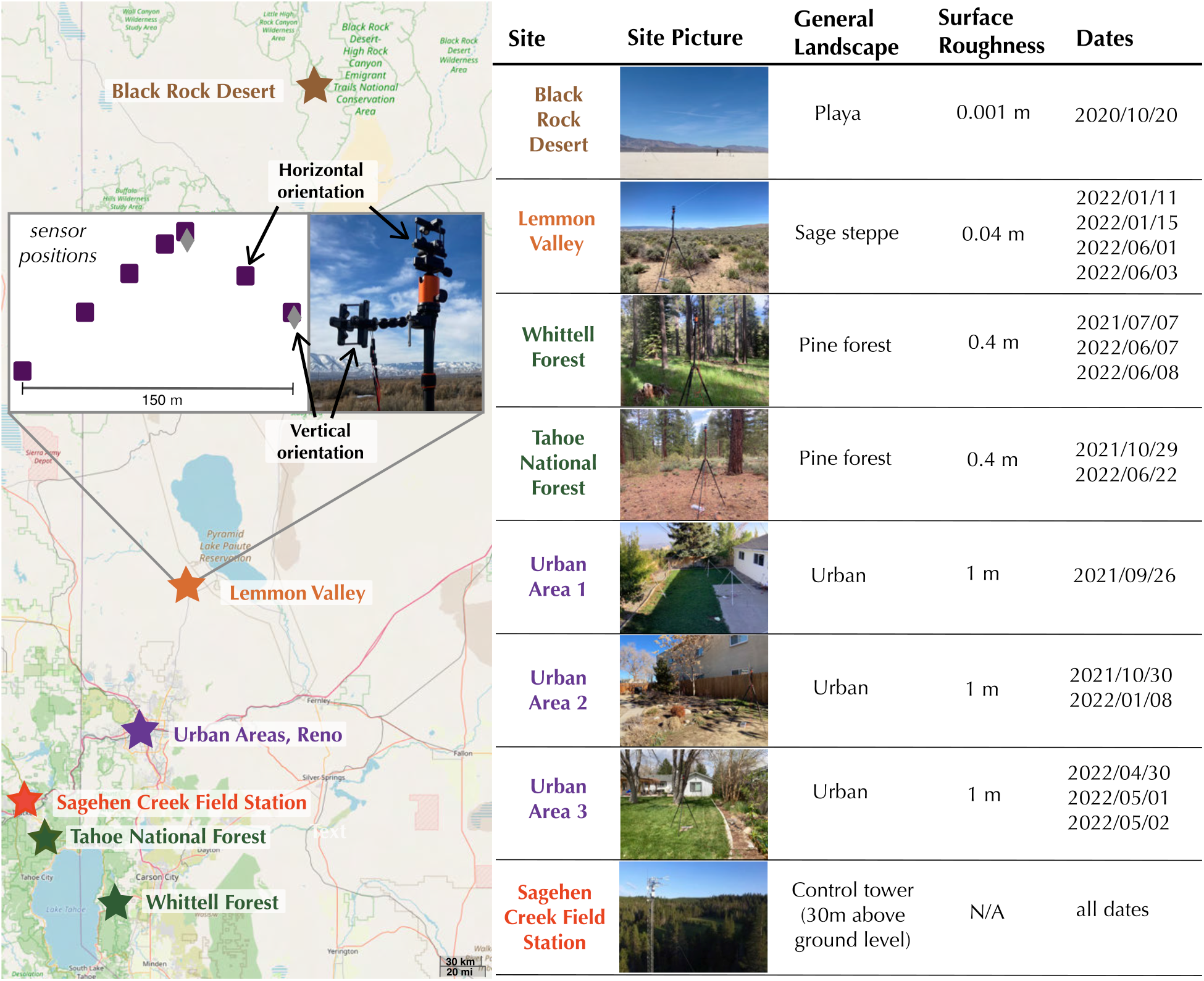
Locations, descriptions and dates of data collection. Sensors were generally oriented in an “L” shape, though this orientation varied slightly across data collections. Each environment will be referred to as its general landscape in subsequent figures. Surface roughness values were estimated based on prior literature [14, 32]. Sagehen Creek Field Station photo image credit to Sebastian Wolf, 06/07/2018.

Both of the forest sites can be characterized as mature coniferous with a mean tree height of approximately 25-30 meters. The 3 urban sites are located within Reno, Nevada. All 3 sites were enclosed by fences approximately 2 meters in height. The fences surrounding urban sites 1 and 2 were wooden whereas urban site 3 had a mix of chain-link and wooden fencing. The surrounding buildings for all urban sites were approximately 6-12 meters tall. Representative pictures are included for each environment in Figure 1. Additionally, Supplemental Figures S1 & S2 show representative horizontal wind speed, horizontal wind direction, and vertical velocity time series for each data collection. Together, these figures demonstrate a diversity of wind speed and direction conditions recorded across days and locations.

### 2.2 Instrumentation and setup

Wind recordings were measured with 3D ultrasonic anemometers (Trisonica™Mini, Anemoment, Longmont, CO) at a rate of 10Hz. The Trisonica™Mini has a specified accuracy of ±0.2 m/s in wind speeds between 0-10 m/s, ±0.2% m/s in wind speeds between 11-30 m/s, and ±1° for wind direction. The anemometers were arranged on tripods at approximately 2 meters above ground level and at distances spanning from 1 meter to 200 meters apart. The spatial layout of sensors varied slightly across data collections based on site constraints. Sensors were generally oriented in an ”L” shape in all data collections aside from the playa, where sensors were arranged in a 3×3 grid. The ”L” shape configuration allowed for more variation in distance between sensors while still encapsulating potential differences caused by landscape heterogeneity. These sensors were manually leveled and oriented toward North with compass readings prior to each experiment. Teensy 3.5 controllers with microSD cards were used for data-logging. A separate GPS receiver (GP-20U7 (56 Channel), Sparkfun) recorded time and location information.

In total, 9 anemometers were utilized throughout our collections. Since the Trisonica™Mini’s design obstructs some vertical flow, we placed 1-2 sensors in a vertical orientation to obtain more accurate vertical flow measurements (sensor orientations shown in Figure 1). In some instances recording malfunctions, adverse weather conditions, and/or wildlife interactions resulted in loss of recording for certain sensors. In some collections, sensors were set up for multiple days. It is well-known that wind regimes shift between night and daytime [33], therefore, for consistency across locations only daytime data (at least 1 hour after sunrise and 1 hour before sunset) was utilized for this analysis.

To quantitatively distinguish between each distinct environment, we assigned each environment an estimated aerodynamic surface roughness length based on prior literature [14, 32, 28]. Though methods for estimating surface roughness with single level anemometer data have previously been proposed [15, 34], they generally require sensor placement above the roughness elements, which was not possible in our forest and urban experiments.

During data pre-processing, minor time measurement offsets were corrected through interpolation. To verify sensor accuracy and noise levels, we recorded measurements in a laminar flow wind tunnel at three different wind speeds (Supplemental Figure S3). Based on these measurements, we chose two different filtering parameters to smooth wind measurements and ran our analysis using both the filtered and unfiltered data. Filtering did not have a strong effect on our results, therefore, all analysis included in this paper was conducted on the unfiltered data. More information about the filtering parameters used can be found in Supplemental Figures S3 & S4.

### 2.3 Statistics: multiple linear regression analysis

To quantitatively compare the effects of key variables that might impact wind speed and wind direction, we conducted multiple linear regression (MLR) using general least squares. In several cases, model diagnostic plots indicated that residual standard errors varied between data collections. To account for this heteroskedasticity and potential correlation within group observations, we implemented cluster-robust standard errors by assigning a group number to each data collection. Regression covariates were z-score normalized to improve interpretation of their relative effect size within each MLR model. To verify that independent variables were not strongly correlated within a model, which would impact both the coefficient estimates and corresponding p-values, Variance Inflation Factor (VIF) was used to assess the covariate structure of our data. All covariates had VIFs between 1-3, indicating that there was no significant multicollinearity present. Statistical diagnostic plots for each model are included in Supplemental Figure S5.

### 2.4 Spatial comparisons: changes in wind characteristics as a function of distance

To visualize the variation of wind direction and speed over distance, we computed the difference in direction and speed at all points in time between each permutation of two sensor pairs. These differences were averaged over discrete 10-minute intervals. We then calculated the distance between sensors using the recorded GPS coordinates. To ensure that our GPS recordings were sufficiently accurate, we manually measured the distance between sensors on several experiments and compared those measurements to the GPS readings. The recorded GPS measurements were always within 2-3 meters of our manual measurements, which is in agreement with the accuracy specified by the GPS manufacturing company and did not have any impact on the interpretation of our results. Since some data was collected on privately owned land, latitude and longitude information was converted to Cartesian coordinates in preparation for making the data publicly available.

To control for the fact that data was collected on different days, in which atmospheric stability and large scale weather conditions may play a role, we used wind speed and direction data from a nearby weather tower (Sagehen Creek Station, CA) during the dates and times of each of our data collections as a variable in our spatial regression analysis (general location shown in Figure 1). This data is managed by the Western Regional Climate Center (WRCC) and is publicly available on their website. The automated weather station is positioned at a height of 30 meters and records the mean wind speed and standard deviation in wind direction over 10-minute intervals. The distances between this tower and our data collections varied from approximately 10 kilometers to 55 kilometers, with exception of the Black Rock Desert, which is approximately 225 kilometers from Sagehen Creek Field Station.

### 2.5 Temporal comparisons: changes in wind characteristics over time

To simplify interpretation of covariate effects across data collections, MLR was chosen for temporal analysis rather than other available time series analysis methods. To do this, we analyzed how the wind direction and wind speed from each data collection was changing over time by computing the standard deviation and mean values of speed and direction over time windows ranging from 0 to 10 minutes at randomly chosen points throughout each dataset.

Since the standard deviation in wind speed values has no upper limit, we normalized these calculations using the mean horizontal wind speed, giving a non-dimensional value which is commonly referred to as turbulence intensity [35], and can be defined as

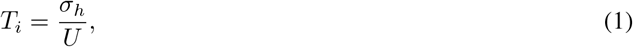

where *σ*_*h*_ is the standard deviation of horizontal wind speed and U is the mean horizontal wind direction.

We implemented similar methods when conducting MLR analysis on the vertical velocity measured by our vertically oriented sensors. To standardize values across datasets, we used vertical turbulence intensity, defined as

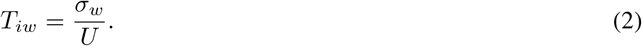

This quantity has been used as a proxy for atmospheric stability and as an estimate for vertical shear in prior field studies seeking to quantify wind profiles in the lower atmospheric boundary layer [25, 26].

In addition to MLR, we investigated temporal variation across environments using both autocorrelation and frequency domain analysis. These methods, however, were not particularly informative of environment driven differences on the timescale of 0-10 minutes. Supplemental Figure S6 shows the mean power spectral densities of horizontal wind speed and direction for each data collection using discrete 10 minute intervals. In most cases, the power spectra were similar, with exception of several days in which wind direction was constant for the majority of sampling time.

The shortcomings of these analyses motivated further visual exploration of the time driven changes in wind direction, eventually leading to the use of probability density distributions. By calculating the standard deviation in direction over time windows ranging from 10^*−*1^ to 10^3^ seconds for all points throughout each data collection, we were able to construct density heatmaps which show the relationship of the standard deviation of wind direction over different time windows for a given time series.

### 2.4 Wind Direction Computations

All calculations involving wind direction were adjusted to account for circularity. When finding the difference in direction between sensors, we used the smaller angle such that the difference was always bounded between [0-180°]. The circular mean was computed using the equation

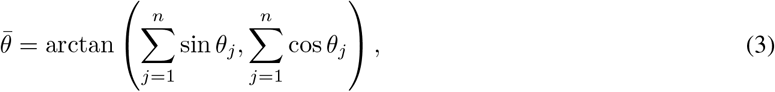

where 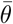 represents the mean angle. The circular standard deviation in direction was calculated as

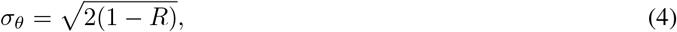

such that *σ*_*θ*_ *∈* [0, *π/*2], where

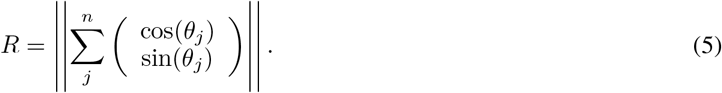

These relations are well documented by [36], [37], and [38].

## 3 Results

### 3.1 Variability of wind speed and direction across space

To understand spatially-driven near-surface wind variation, we compared the absolute difference in horizontal wind direction and speed between pairs of sensors over discrete 10 minute periods. Figure 2A demonstrates this differencing method for two sensors over a time period of 50 minutes. By repeating this calculation for all permutations of two sensor pairs across each data collection, we recovered statistics which provide intuition on how different wind observations were for a given time period over distances ranging from 0-200 meters. Table 1A details MLR results using the data shown in Figure 2A where the 10-minute mean difference in direction between sensors is the dependent variable and the independent variables are the 10-minute mean wind speed, estimated environment surface roughness, and distance between sensors. Similarly, Table 1B details MLR results using the data shown in Figure 2B where the 10-minute mean difference in wind speed between sensors is the response variable. The following subsections detail relationships observed as a function of these wind direction and speed differences.

**Table 1:** Multiple linear regression coefficients, standard errors, and p-values for A) absolute difference in wind direction between sensors (adjusted *R*^2^ = 0.662) and B) absolute difference in wind speed between sensors (*R*^2^ = 0.229). Estimated surface roughness was used as a proxy for environment as a means of quantifying surface complexity. The 10-minute standard deviations in wind direction and 10-minute mean wind speeds measured by Sagehen Creek Field Station were used as a control in regressions A and B, respectively.

| (a) | Variable | Normalized Coefficient | Standard Error | P-value | (b) | Variable | Normalized Coefficient | Standard Error | P-value |
| --- | --- | --- | --- | --- | --- | --- | --- | --- | --- |
|  | 10-min mean wind speed | -0.7318 | 0.093 | <0.001 |  | 10-min mean wind speed | 0.8563 | 0.207 | <0.001 |
|  | 10-min mean wind speed x Environment | -0.3207 | 0.073 | <0.001 |  | 10-min mean wind speed x Environment | 0.5491 | 0.157 | <0.001 |
|  | Environment | 0.3027 | 0.094 | 0.001 |  | Environment | 0.3326 | 0.123 | 0.007 |
|  | Intercept | -0.2999 | 0.140 | 0.032 |  | Intercept | 0.2826 | 0.125 | 0.024 |
|  | Distance x Environment | -0.2364 | 0.139 | 0.089 |  | Distance x Environment | 0.0529 | 0.067 | 0.459 |
|  | Control x Environment | 0.1412 | 0.036 | <0.001 |  | Distance | 0.0511 | 0.069 | 0.459 |
|  | Control | -0.0717 | 0.045 | 0.108 |  | Control | -0.0305 | 0.055 | 0.580 |
|  | Distance | 0.0712 | 0.163 | 0.662 |  | Control x Environment | 0.0044 | 0.055 | 0.936 |

**Figure 2:**
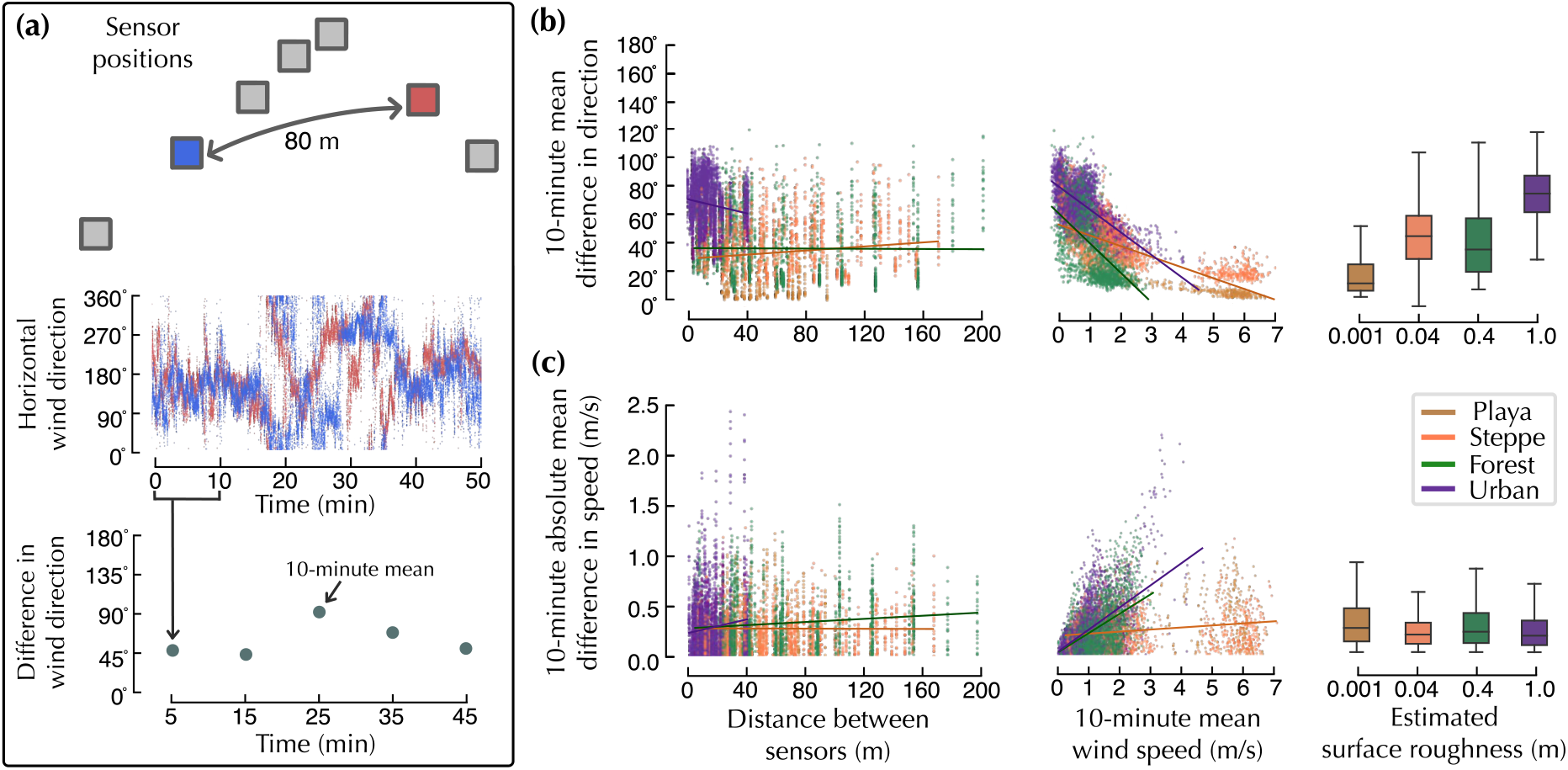
Spatial variability in wind direction is primarily correlated with wind speed and environment surface complexity. A) Visual demonstration of difference in direction calculations between two signals recorded at a distance of 80 meters apart. B) Summary statistics showing the 10-minute mean difference in wind direction between sensors as a function of distance, wind speed, and estimated surface roughness. C) Summary statistics showing the 10-minute mean difference in wind speed between sensors as a function of distance, wind speed, and estimated surface roughness.

### Directional variability decreases with increasing mean wind speed

From our regression analysis, we found the difference in wind direction between sensors to be most strongly correlated with mean wind speed (normalized coefficient = -0.7318, p-value *<*0.001, Table 1A). Across all data collections, the difference in wind direction was highest when mean wind speeds were less than 2 m/s, with variability dropping significantly around the 3 m/s threshold (Fig. 2B). This phenomenon is consistent with prior atmospheric science literature indicating that wind direction variability is highest in low wind speed conditions [19, 22, 39].

It should be noted that our MLR model contained some residual heteroskedasticity (Supplemental Figure S5). While running diagnostics, we found that the majority of heteroskedasticity was due to the transitional relationship between mean wind speed and wind direction which occurs when wind speeds increase beyond approximately 2 m/s. Implementing the same regression analysis on a subset of the data in which mean wind speeds were constrained to under 2 m/s corrected for the majority of residual heteroskedasticity. This correction did not strongly impact regression standard errors due to use of cluster-robust covariance, however, the normalized coefficient for mean wind speed dropped to -0.4943 (p-value *<*0.001). This indicates that mean wind speed may become a more dominate influence of wind direction variability above the 2 m/s threshold.

### Environments with higher surface complexity exhibit larger directional variability

Environment complexity (quantified by using estimated surface roughness) also corresponded with greater differences in direction measurements between sensors (normalized coefficient = 0.3027, p-value = 0.001). There was a clear difference in directional changes observed between the playa and urban environments, whereas directional differences observed in the sage steppe and forest environments had more similar distributions (Fig. 2B). Interestingly, the mean difference in direction between sensors was higher in sage steppe (43.90°) than in forest (38.86°). Even still, the observed range of minimum and maximum differences was lower in sage steppe, and the spread of the interquartile range was smaller (26.57° versus 32.92° for sage steppe and forest, respectively).

### The distance between sensors did not greatly influence observed directional differences

The 10-minute mean difference in wind direction across sensors did not significantly depend on distance alone (p-value = 0.662) or on interactions between distance and environment (p-value = 0.089). Figure 2B shows a visible upward trend of approximately 5° over 200 meters in forest and playa environments, although the range of differences observed between sensors still varied as widely as 20° to 100° across all distances. Though there is no strong linear relationship between difference in direction and distance between sensors, Figure 2B hints at a non-linear relationship in the forest and steppe environments in distances greater than 100 meters.

Counter-intuitively, Figure 2B shows a downward trend in directional changes between sensors in urban environments of approximately 10°. This could be due to highly obstructed flow near fences and buildings, which may create turbulent flow in the immediate surroundings but have some uniformity in middling areas where the space is more open. Since the urban experiments were conducted on privately owned land, our spatial distribution of sensors was limited to areas less than 50m^2^. It is possible the observed downward trend would even out in larger urban areas such as city parks.

### The effects of control data were minimal

The control data from Sagehen Creek Field Station (10-minute standard deviation in direction observed at 30 meters above ground level) was not significant as a stand-alone predictor for difference in direction between sensors (p-value = 0.108, Table 1A). This indicates that collecting data on different days did not strongly influence MLR results. We note that the interaction term between our control data and estimated surface roughness was significant, though the coefficient itself was relatively small (normalized coefficient = 0.1412, p-value *<*0.001). This indicates that a portion of the directional variability observed across environments may be attributed to the difference in locations that data was collected.

### The absolute difference in speed between sensors showed no strong correlations

Figure 2C demonstrates that the absolute difference in horizontal wind speed between sensors is not well described as a function of distance, mean wind speed, or environment. From MLR model results, we find that although several of the terms have significant p-values, the adjusted *R*^2^ is quite low (*R*^2^= 0.229, Table 1B), suggesting that this model does not explain much of the variability. Further, the residual diagnostic plots in Supplemental Figure S5 indicated that this model had a particularly high level of heteroskedasticity and outliers.

To assess the impacts of these residual issues on model estimates and standard errors, we performed a Box Cox transformation (*λ* = 0.326) and removed outlier data. This corrected for most of the heteroskedasticity and resulted in an adjusted *R*^2^ of 0.153, along with an approximate 10% reduction in normalized coefficient values for environment surface roughness and 10-minute mean wind speed. Though we include the MLR results (Table 1B) and residual diagnostic plots (Supplemental Figure S5) of the original model for ease of interpretation, it should be noted that the reduction in *R*^2^ after transformation indicates that the difference in speed between sensors is not well predicted by distance, environment, or mean wind speed.

### 3.2 Variability of wind speed and direction over time

We initially investigated temporal driven variations in wind speed and direction using autocorrelation and power spectral density analyses. Since temporal changes can be affected by both mean wind speed and environment, decoupling and quantifying the relative effects of each variable over small time windows proved challenging when using these methods. Supplemental Figure S6 shows the 10-minute averaged power spectral density for each day of data collected. Though spectra characteristics were relatively indistinguishable when directly comparing environments, it can be noted that the -5/3 Kolmogorov spectrum, which is often used to describe turbulence [40], roughly holds true in the frequency ranges between 10^*−*1^ and 10^1^ Hz for most days and environments.

The exploration of these two analyses made way for a sliding time window approach, which is illustrated in Figure 3A. Figure 3B shows three distinct wind direction conditions (consistent, large infrequent shifts, and constant fluctuations) and Figure 3C displays the corresponding density heatmaps obtained by computing the standard deviation in wind direction over time windows ranging from 10^*−*1^ to 10^3^ for all points throughout the given time series. It can be seen in all scenarios that there is a positive increase in *σ*_*θ*_ as the time window increases. In cases of consistent wind direction, *σ*_*θ*_ levels off within 1-2 seconds, whereas we see *σ*_*θ*_ level off at around 100 seconds in the most variable case. These three examples are included to provide intuition on the time driven variations in wind direction and the typical time window length at which *σ*_*θ*_ levels out in different conditions.

**Figure 3:**
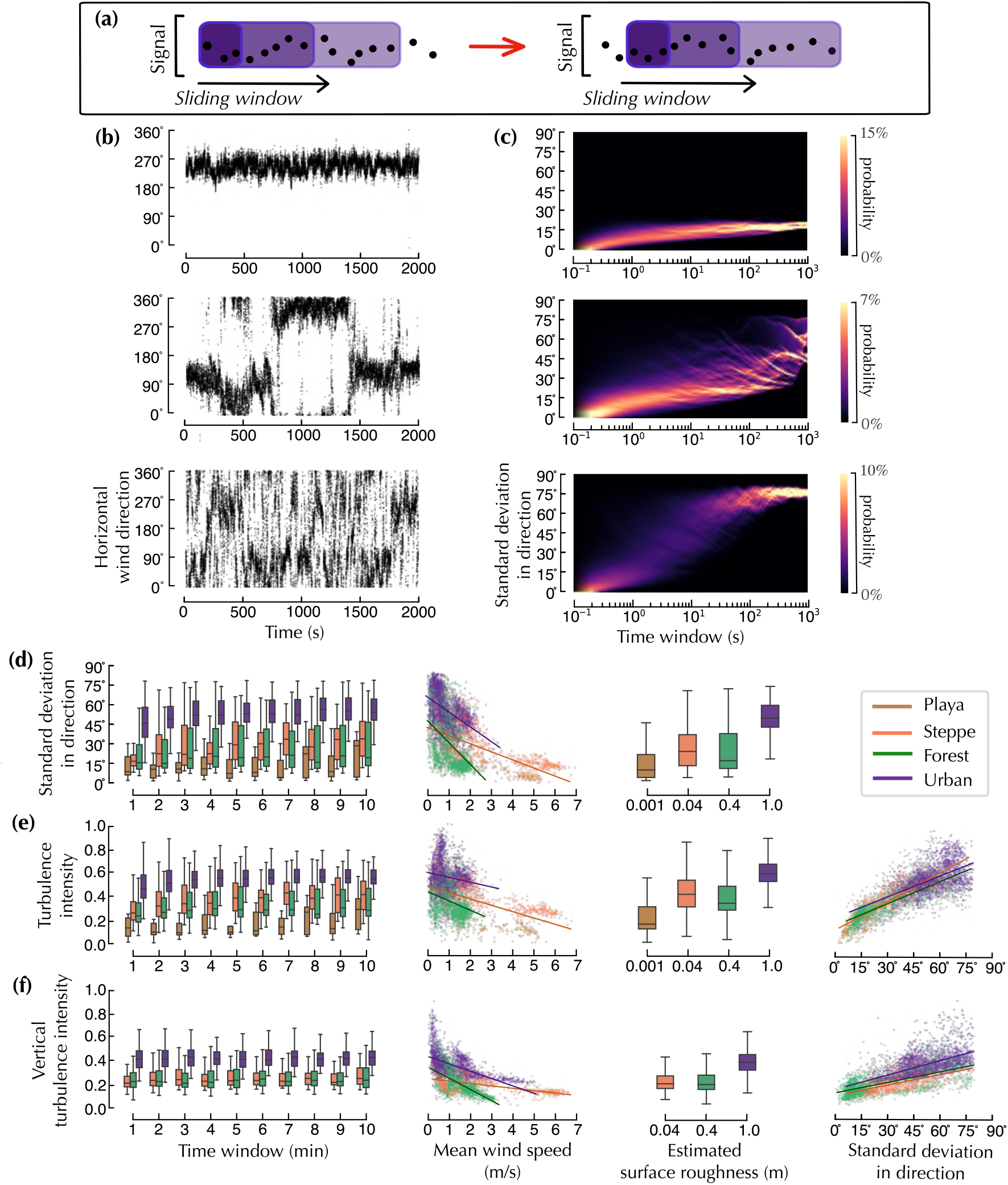
Temporal variations in wind statistics can be captured by using varying time window lengths across the observed signal. A) Illustration of sliding window calculations used to compute the temporal variations in *σ*_*θ*_, *T*_*i*_, and *T*_*iw*_ for each dataset. B) Three examples of wind direction over time: constant, large infrequent shifts, and highly variable. C) Standard deviation in wind direction over varying time windows from 0 to 1000 seconds, calculated using the method described by A. D) Summary statistics comparing the standard deviation in direction across environments over linearly spaced time intervals ranging from 0 to 10 minutes. E) Summary statistics comparing the turbulence intensity across environments over linearly spaced time intervals ranging from 0 to 10 minutes. F) Summary statistics comparing the vertical turbulence intensity across environments over linearly space timed intervals ranging from 0 to 10 minutes. Note that we did not have any vertically oriented sensors in the playa and thus vertical turbulence intensity values from that data are absent.

### Standard deviations in direction increase with increasing time windows

As can be seen in Figure 3D, the standard deviation in direction (*σ*_*θ*_) generally increased over time in all environments (normalized coefficient = 0.1809, p-value *<*0.001), though the relative effect size indicates that time window is less important than mean wind speed and environment surface roughness (Table 2). The temporal variations of each data collection can be seen with more detail in Figure 4, which is organized as a function of overall mean horizontal wind speed and environment.

**Table 2:** Multiple linear regression coefficients, standard errors, and p-values for A) standard deviation in wind direction (adjusted *R*^2^ = 0.519), B) turbulence intensity (adjusted *R*^2^ = 0.714), and C) vertical turbulence intensity (adjusted *R*^2^ = 0.610). Estimated surface roughness is used as a proxy for environment. Corresponding residual diagnostic plots are included in Supplemental Figure S5.

| (a) Variable | Normalized Coefficient | Standard Error | P-value | (b) Variable | Normalized Coefficient | Standard Error | P-value | (c) Variable | Normalized Coefficient | Standard Error | P-value |
| --- | --- | --- | --- | --- | --- | --- | --- | --- | --- | --- | --- |
| Mean wind speed | -0.5640 | 0.141 | <0.001 | Std. deviation in direction | 0.7667 | 0.072 | <0.001 | Std. deviation in direction | 0.5898 | 0.063 | <0.001 |
| Environment | 0.4068 | 0.114 | <0.001 | Environment | 0.0988 | 0.050 | 0.049 | Environment | 0.3115 | 0.086 | <0.001 |
| Mean wind speed x Environment | -0.2365 | 0.121 | 0.050 | Time window | 0.0562 | 0.013 | <0.001 | Environment x Std. deviation in direction | 0.0967 | 0.077 | 0.207 |
| Time window | 0.1809 | 0.028 | <0.001 | Environment x Std. deviation in direction | -0.0501 | 0.085 | 0.556 | Intercept | -0.0397 | 0.098 | 0.685 |
| Intercept | -0.1067 | 0.179 | 0.550 | Intercept | 0.0358 | 0.049 | 0.464 | Time window | -0.0544 | 0.019 | 0.004 |
| Mean wind speed x Time window | -0.0440 | 0.012 | <0.001 | Time window x Std. deviation in direction | -0.0293 | 0.014 | 0.037 | Time window x Std. deviation in direction | -0.0339 | 0.018 | 0.053 |

**Figure 4:**
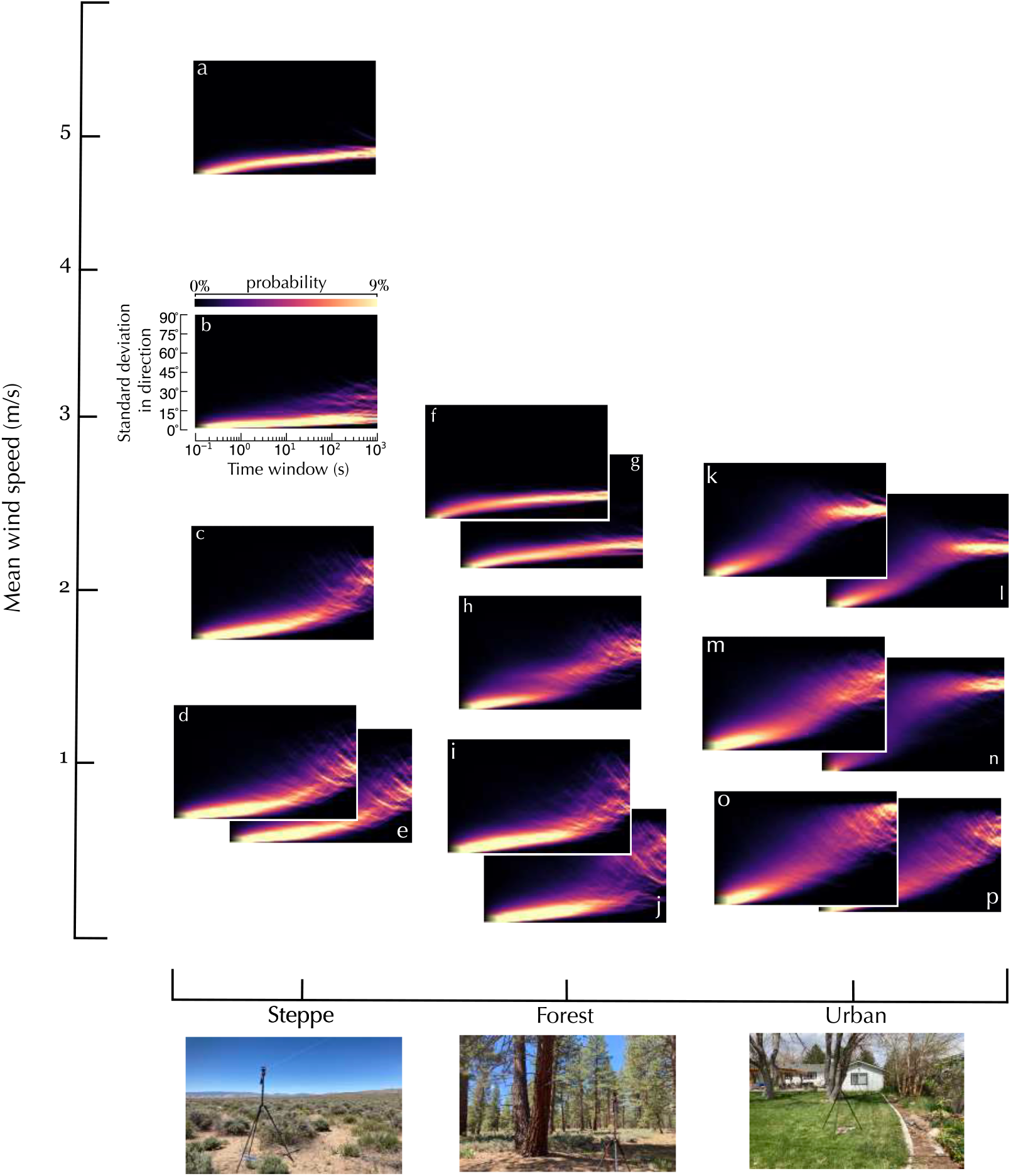
The standard deviation in wind direction generally increases for larger time windows and is a function of both wind speed and environment. We computed the standard deviation in wind direction over varying time window lengths for all points throughout each data collection. Each subplot (a-p) represents the probability distribution corresponding to a time series, which can be found in Supplemental Figures S1 & S2. Note that subplot b corresponds to data collected in the playa, but is grouped with the steppe environment for visual simplicity.

**Figure 5:**
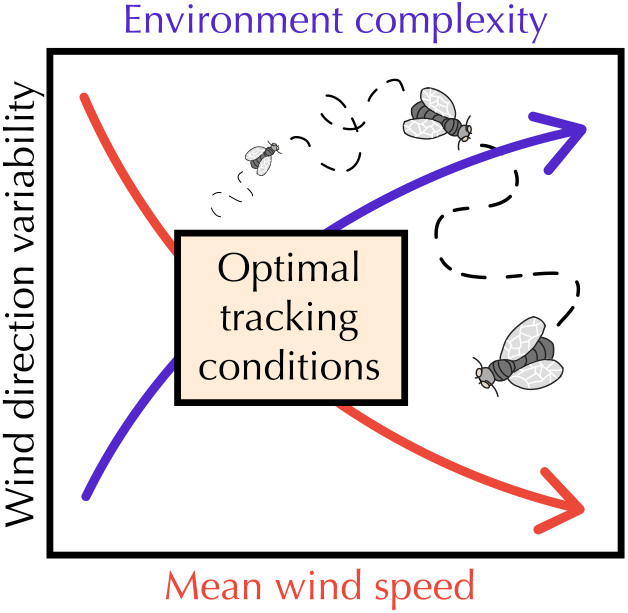
Optimal long distance plume tracking conditions are likely found at intermediate wind speeds and environmental complexities.

### Standard deviations in direction are largely a function of environment and mean wind speed

From MLR results, we found that *σ*_*θ*_ was negatively correlated with mean wind speed (normalized coefficient = -0.5640, p-value *<*0.001). Figure 3D shows that the relationship between *σ*_*θ*_ and mean wind speed was similar to the directional variability observed in our spatial analysis (Figure 2B). Furthermore, Figure 4 demonstrates that the maximum value of *σ*_*θ*_ decreased with increasing mean wind speed across environments.

Environment surface complexity also played a sizable role in temporal variations of *σ*_*θ*_ (normalized coefficient = 0.4068, p-value *<*0.001). It should be noted that although we collected wind data over a variety of days with differing conditions, the overall mean horizontal wind speed in forest and urban environments never exceeded 3 m/s. These observations indicate that near-surface mean wind flow may generally be lower in environments with more surface roughness elements.

Our results are consistent with prior temporal analysis of surface wind patterns in the nocturnal stable boundary layer [24, 22]. [22] reported values of near-surface *σ*_*θ*_ around 50° in nocturnal conditions under low wind speeds. Mahrt also found that *σ*_*θ*_ decreases substantially around the 2 m/s wind speed threshold. Although Mahrt primarily focused on nocturnal wind regimes, he did note that the standard deviation in direction would typically be higher in day time regimes due to convective processes, as well as in more complex environments. These ranges are in agreement with our findings, though we see some higher values of *σ*_*θ*_ (up to approximately 75°) in the urban data.

### Horizontal and vertical turbulence intensity are not impacted by time window length

Table 2B shows that although the relationship between the horizontal turbulent intensity (*T*_*i*_) and time window in our MLR model was statistically significant (p-value *<*0.001), the normalized coefficient is very small (0.0562). The effect size of time window is equally small with respect to the vertical turbulent intensity (*T*_*iw*_) (normalized coefficient = -0.0544, p-value = 0.004, Table 2C). These results indicate that time window size has very little effect on *T*_*i*_ and *T*_*iw*_ over the scale of 0-10 minutes. In addition to Figures 3E & F, direct relationships between horizontal and vertical wind speed are included in Supplemental Figure S7, and demonstrate a general correlation between mean speeds and standard deviations in speed across environments.

### The role of environment on horizontal and vertical turbulence intensity

Figures 3E & F demonstrate that values of *T*_*i*_ and *T*_*iw*_ were generally larger in environments with higher estimated surface roughness. However, MLR results in Tables 2B & C indicate that the relative effects of environment surface complexity appear to be much stronger with respect to *T*_*iw*_ (normalized coefficient = 0.3115, p-value *<*0.001) than with *T*_*i*_ (normalized coefficient = 0.0988, p-value = 0.049). Overall, values of *T*_*i*_ were higher in the sage steppe environment than in the forest environment, whereas values of *T*_*iw*_ were much closer in the steppe and forest environments. It is worth noting that these wind statistics were taken at a height of 2 meters above ground level, and would likely be different for winds observed at canopy top in forest environments.

Several prior studies have been conducted within the urban surface roughness sublayer in an attempt to better understand the complexities of disturbed urban flow [26, 25]. [25] reported values of 0.3 for *T*_*iw*_ in 3 different cities. Here, we report values closer to 0.4 for our urban data. Though these differences could be due to atmospheric stability on the days we collected data, [41] notes that the overall effects of atmospheric stability are expected to be minimal due the strong mechanical mixing of wind forced by surrounding buildings. Thus, we speculate a possible reason for this discrepancy may be the positioning of our vertically oriented sensors, which were typically within 1-2 meters of a fence or building and may have resulted in increased vertical flow due to recirculating winds created by the surrounding fencing and infrastructures.

### Horizontal and vertical turbulence intensity are strongly correlated with standard deviation in direction

From Figures 3E & F and MLR results in Tables 2B & C, it can be seen that both *T*_*i*_ and *T*_*iw*_ are strongly correlated with *σ*_*θ*_. These relationships are logical, given that *T*_*i*_, *T*_*iw*_, and *σ*_*θ*_ are often used as general indicators of near-surface turbulence. This relation may be of particular interest to researchers interested in replicating natural wind conditions in wind tunnels, where it is possible to increase *T*_*i*_ without inducing large values of *σ*_*θ*_ through changes in wind speed.

## 4 Discussion

Our results highlight that near-surface wind variability over spatial and temporal scales relevant to foraging insects is primarily influenced by mean wind speed and environment surface complexity. Consistent with prior literature, we find that wind directional variability is inversely proportional to mean wind speed. We further observe that increases in surface complexity contribute to higher variation in wind direction both temporally and spatially. Notably, there was a strong relationship between turbulence intensity and the standard deviation in direction. Though our results indicated that spatial variation and time scale have some impact on wind variability, these impacts are much smaller than those caused by both environment and wind speed.

### Significance of wind speed on near-surface wind variability

Across environments, wind direction variability decreases with increasing wind speeds. This effect becomes increasingly pronounced when mean speeds surpass the 2 m/s threshold. Though insects can passively travel in higher wind speeds, they must actively navigate through wind when tracking an odor plume. The range of wind speeds in which insects have been observed performing these tasks varies based on the size of the insect. Wind tunnel experiments with moths show that they can successfully navigate mechanically induced turbulent conditions in 1 m/s wind speeds [42], whereas *Drosophila* tracking tasks are typically conducted in wind speeds around 0.4 m/s [43, 44]. Field studies suggest that at wind speeds higher than 1 m/s, *Drosophila* are likely to be substantially advected by the wind [11]. Furthermore, field based studies with bees show that flower foraging rates drop in higher wind speeds [45, 46, 47]. [47] found a 38% decrease in flower visits by bees when wind speeds increased from the range of 0-1 m/s to 2.5-3.5 m/s. Together, these prior studies suggest that many insects are most likely to perform odor plume tracking behaviors in wind speeds of less than 3 m/s.

From our results, we expect that insects tracking odor plumes in the 0-2 m/s wind speed range will also experience higher levels of directional variability, leading to more meandering and disperse plume shapes. [42] reported that moths in wind tunnels performed better with mechanically induced turbulence than in purely laminar flow due to the two to three-fold increase in the plume’s cross sectional area 1m downwind. Though *T*_*i*_ was not quantified in this study, [48] gives more information on the turbulence intensity observed in their setup, with reported values of *T*_*i*_ around 5-20% depending on measurement location within the tunnel. *σ*_*θ*_ was not quantified, but was likely small compared to the values we observe in outdoor environments due to the constraints of a unidirectional wind tunnel. In a recent field study, [49] found that temporal changes in wind speed and turbulence increased the amount of bee visits to fragmented forested sites due to increases in odor dispersion. Both of these results support the idea that insects may be better equipped to extract odor plume information in wind conditions with higher directional variability. In wind tunnels,[3] generating larger values of *T*_*i*_ is generally easier than creating higher fluctuations in *σ*_*θ*_ due fan orientation constraints. Since we see that *T*_*i*_ is strongly correlated with *σ*_*θ*_, the ability to increase *σ*_*θ*_ in future wind tunnel experiments will be important for uncovering insect tracking strategies in more natural wind conditions.

### Significance of surface complexity on near-surface wind variability

Estimated surface roughness had the second largest effect size in both spatial and temporal regressions, indicating that wind direction is largely a function of both wind speed and surface complexity. There was a distinct separation in directional variability and turbulence intensity between the least complex (playa) and most complex (urban) environments. This separation was not as clear when comparing differences between the sage steppe and forest environments. This may, in part, be due to the height of our sensor placement, which was approximately 1 meter above the mean roughness element in the steppe experiments and approximately 20+ meters below the mean roughness element in the forest experiments.

There is evidence to support that wind variability is elevated directly above surface roughness elements. The atmospheric layer just above the roughness elements is often referred to as the “mixing layer” [50, 35]. [25] observed consistently higher values of turbulence quantities “near and above H”, where H is the mean building height. Conversely, [51] noted less wind direction variability within the “bole-space” spanning from the ground to the lower canopy of a conifer forest when compared to wind direction observations above the canopy. From these works, we posit that less variability may have been observed in the steppe environment if wind was measured below the mean roughness element. Future field work aimed toward collecting data in various other environments and geographical locations would help to fill the knowledge gap surrounding near-surface wind variability in environments of middling surface complexity.

It is important to note that some of our analyses were limited due to the use of one sensor height. This impacted our ability to determine the atmospheric stability and more accurately estimate surface roughness values for each data collection. Even still, insects are typically not inhabiting higher air-spaces unless migrating. More information about the atmospheric conditions affecting these migration events can be found in [52] and [53]. Though insects inhabit various levels of air-space, an insect’s flight altitude while tracking an odor plume will generally be dictated by the height from which the odor is dispersed [54]. Since prior works suggest that insects will often travel closer to the ground while tracking an odor plume [30, 3], limiting analysis to the lowest level of the surface boundary layer provides insight on sensory information that an insect would have access to while actively pursuing an odor plume. Though there may be some large scale impacts on the local wind due to non-localized atmospheric instability, topography, and daytime heat convection, both the MLR control estimates in Table 1 and power spectral density analysis in Supplemental Figure S6 indicate that these impacts were not largely discernible on the time scale of 10 minutes.

### How might wind variability relate to insect plume tracking success?

Supplemental Figures S1 & S2 demonstrate the variability of wind speed and direction measurements observed across environments and days. In some cases wind speeds were quite low, with highly variable wind direction. In other cases, wind direction was fairly consistent. Overall, the wind conditions an insect might encounter are quite diverse, suggesting that insects may implement different odor tracking strategies based on prevailing flow conditions. Additionally, the time and length scales over which canonical cast-and-surge plume tracking can occur may be impacted by the wind variability an insect encounters. In open spaces, such as playa and sage steppe, overall mean wind speeds can often reach higher values than in environments with greater obstructions. In these scenarios, insects may be utilizing the higher wind speeds and less variable wind direction to track plumes over larger distances than they would in more spatially complex areas. Though our findings suggest that wind direction becomes less variable with increasing speed, prior studies indicate that insects can struggle to fly in high wind speeds. Taken together, we hypothesize that there may be an optimal range of conditions in which wind speeds do not present significant bio-mechanical disadvantages and environment surface complexity does not greatly obstruct wind direction, thus providing the optimal ratio of foraging options and wind conditions for successful long distance odor plume tracking.

Insect populations have seen major declines over the past century, with habitat loss due to agriculture and urbanization conversion being a key driver [55, 56]. Farmland wind erosion has been a growing concern over the past century, and is caused by strong winds over bare soils [57, 58]. Conversely, urbanization drastically increases surface roughness and creates strong disturbances in natural wind flow. These types of land developments drastically alter environment surface roughness and wind flow in ways that are likely to impede an insect’s ability to successfully track odor plumes over long distances.

### Implications for future plume tracking experiments

Although spatially driven directional variability was assessed by finding the observed difference in direction recorded at two locations, whereas temporal variability was assessed using the standard deviation over varying time windows, our results show that the corresponding relationships between environment and wind speed are remarkably similar. This indicates that the use of temporal statistics with one sensor may be sufficient for future studies aimed at further quantification of environment specific differences. That said, the spatially driven non-linear relationship observed in directional variability over distances between 100-200 meters in the forest and steppe environments (Figure 2B may warrant future studies with similar sensor setups.

Our knowledge of how insects successfully track odor plumes in outdoor spaces remains limited due to the underlying complexities of near-surface wind dynamics, which influence odor plume dispersion and insect flight strategies. The observed near-surface wind conditions presented in this study demonstrate significant diversity of wind conditions across environments and days of sampling. In many cases, the standard deviation in wind direction ranged from 15° to as much as 75° on the time scale of 1-10 minutes. Under such conditions odor plumes will be more dispersed and meandering, and the canonical cast-and-surge behavior that insects employ in laminar wind conditions [59, 3] is unlikely to be effective. Instead, insects may need to rely on a time history of odor and wind information [43, 60, 61], or a different strategy altogether. Discovering what these strategies are will require controlled laboratory settings in novel wind tunnel assays where directional variability can be controlled to match the linear relationship with turbulent intensity that we describe in Figure 3E-F.

## 5 Acknowledgements

We thank Stephen Drake for discussions relating our work to atmospheric sciences and comments on the manuscript, and Paul Hurtado for statistical analysis advice, and Benjamin Cellini for comments on the manuscript. We thank Hunter Noble and Sarah Bisbing for help with access and permits to collect data at the Whittell Forest and Wildlife Area, the Black Rock Field Office of the Bureau of Land Management for permitting our use of the Black Rock Desert, and various private land owners for permission to record wind data on their properties.

## 6 Funding Statement

This work was partially supported by grants from the Air Force Research Laboratory to F.v.B. (FA8651-20-1-0002), Air Force Office of Scientific Research to F.v.B. (FA9550-21-0122), the Sloan Foundation to F.v.B. (FG-2020-13422), the National Science Foundation AI Institute in Dynamic Systems (2112085), and an NSF EPSCOR UROP Scholarship and Nevada DRIVE Fellowship to J.H.

## 7 Conflict of interest

The authors declare no conflict of interest.

## 8 Data Availability Statement

Data will be made available on Figshare on publication. Code used for data analysis will be made available on Github on publication.

## S1 Supporting Information

Additional figures are included as supplementary material for the reader’s reference.

**Figure S1:**
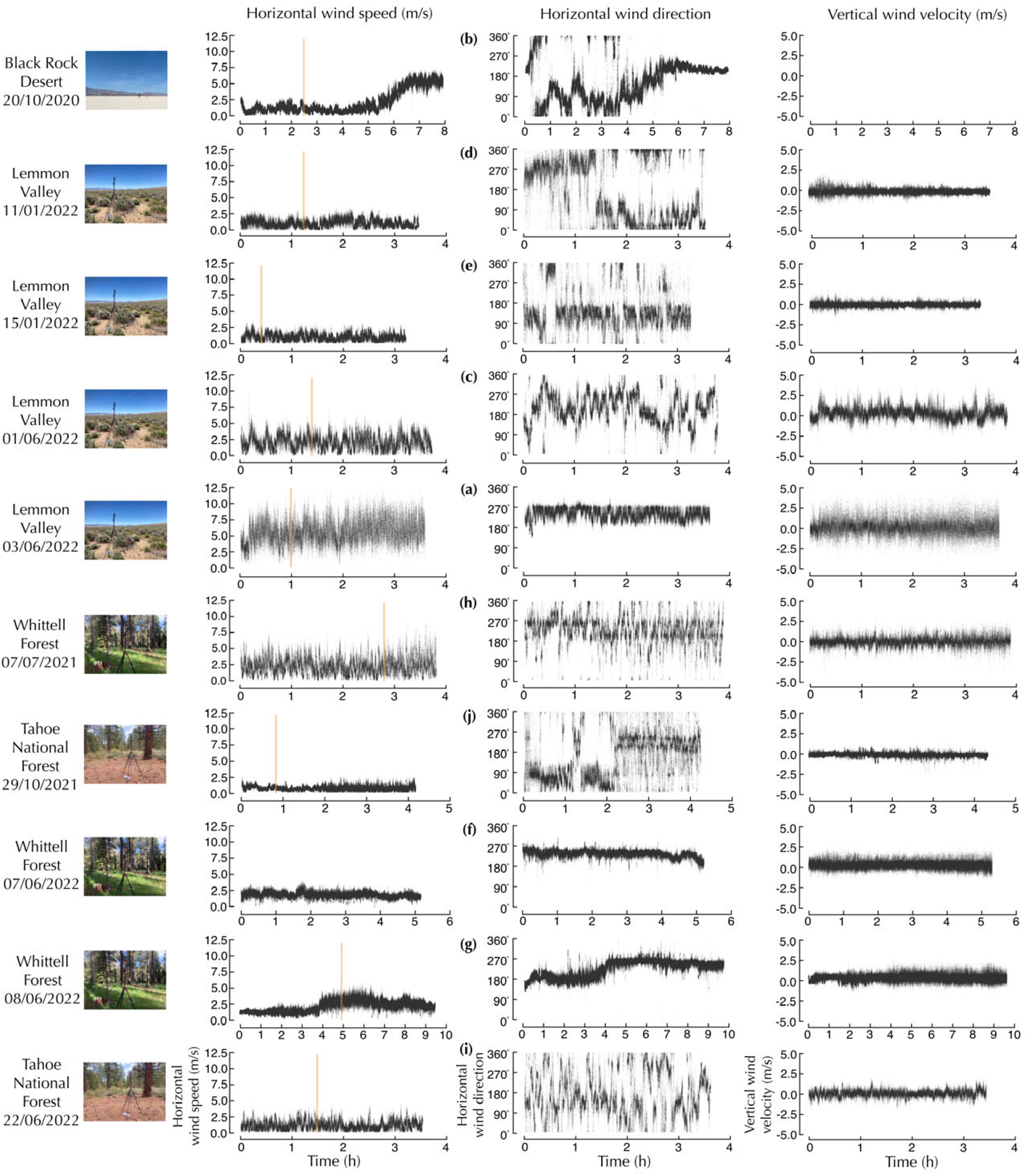
From left to right: horizontal wind speed, horizontal wind direction and vertical velocity. Each row corresponds to a data collection organized from oldest (2020) to most recent (2022). The orange lines represent approximate solar noon based on date and location of data collection. The letters next to wind direction correspond to heatmap plots in Figure 4.

**Figure S2:**
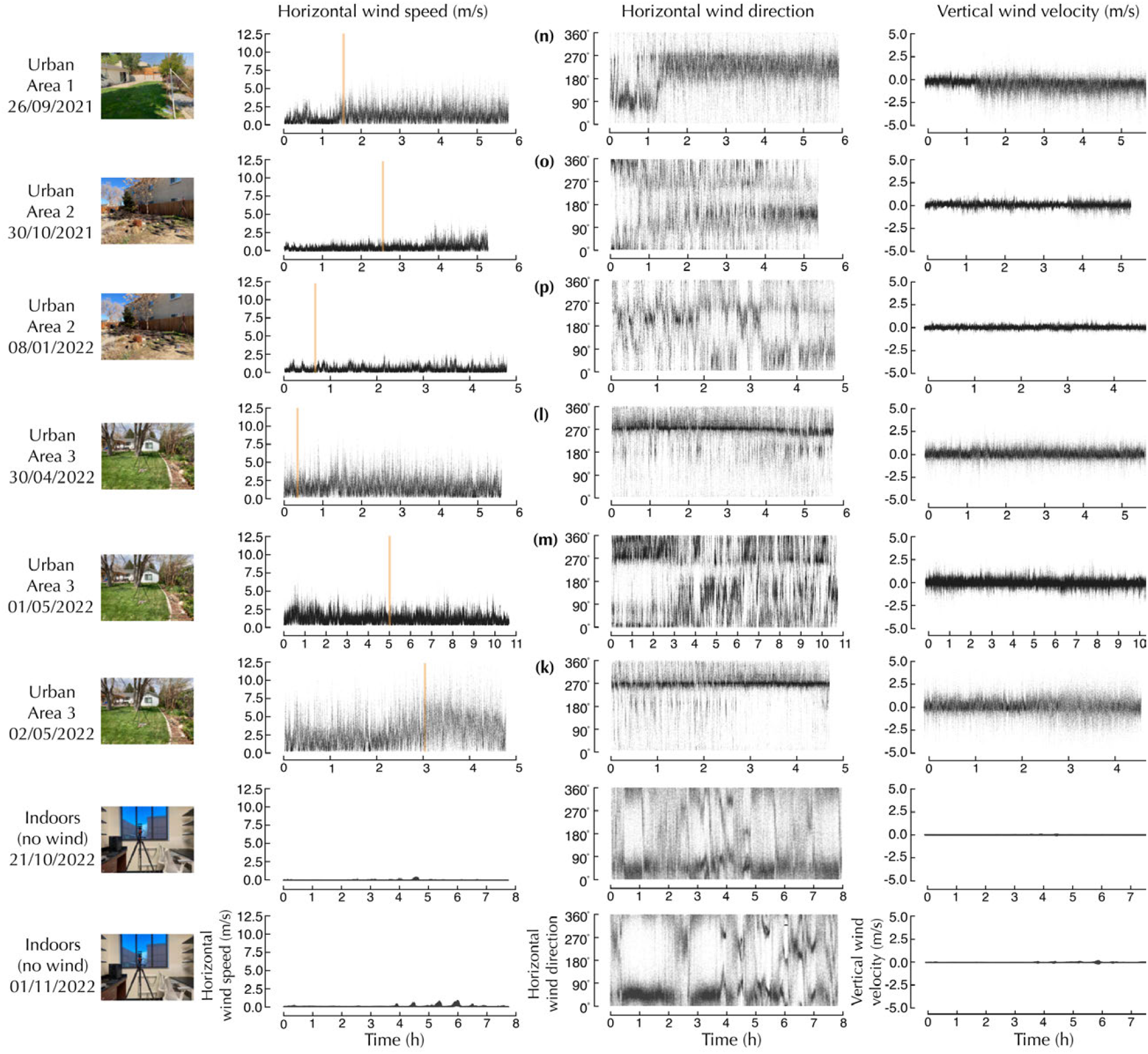
From left to right: horizontal wind speed, horizontal wind direction and vertical velocity. Included are two additional signals recorded indoors overnight. The orange lines represent approximate solar noon based on date and location of data collection. The letters next to wind direction correspond to heatmap plots in Figure 4.

**Figure S3:**
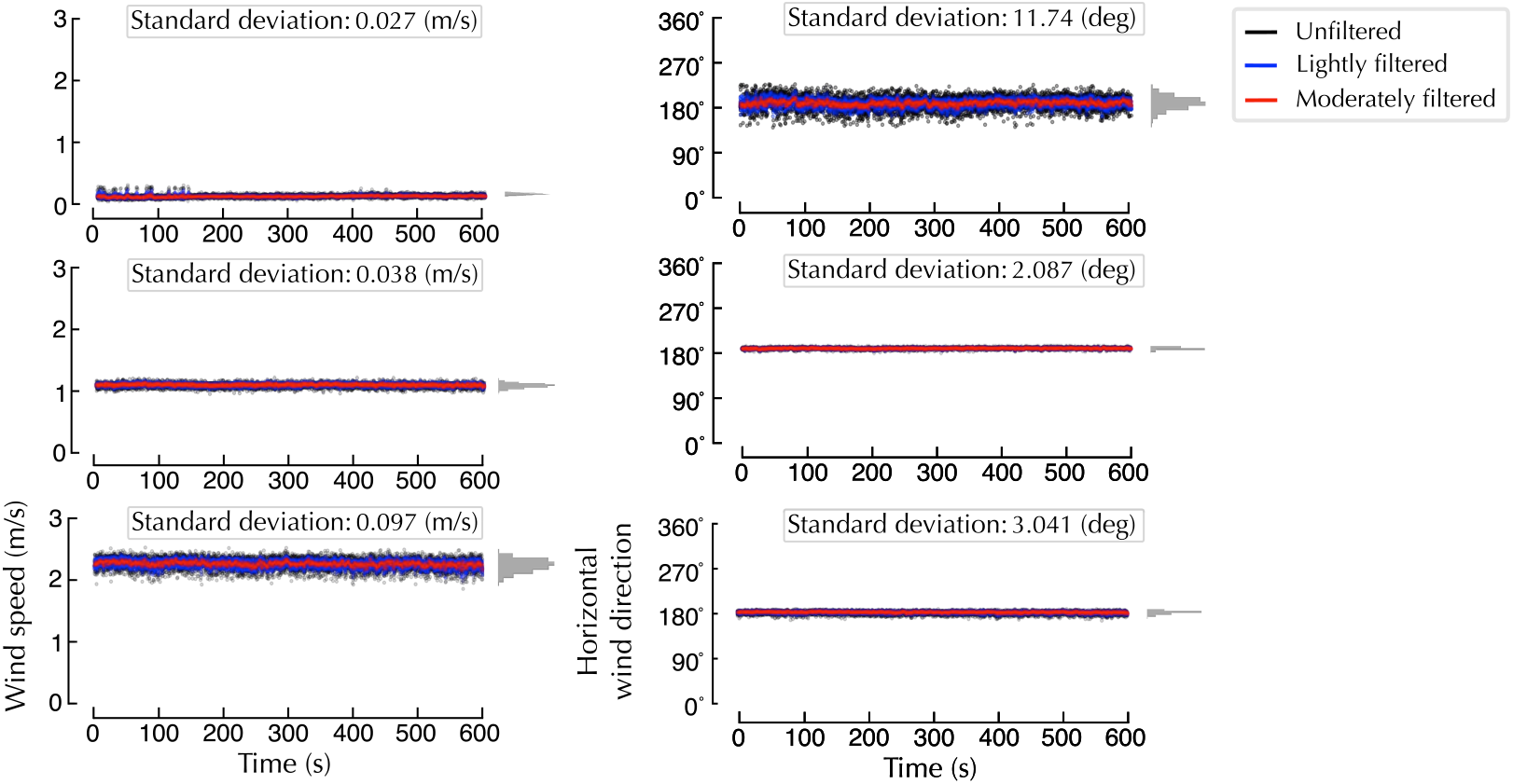
Noise test for Trisonica™Mini 3D ultrasonic anemometer. Wind data was collected in a wind tunnel at 3 different wind speeds. A 3rd order low-pass Butterworth filter was used for smoothing, with cutoff frequencies of 0.5 Hz and 0.1 Hz for the light and moderate filtering, respectively. The standard deviations and histogram distributions correspond to the unfiltered data.

**Figure S4:**
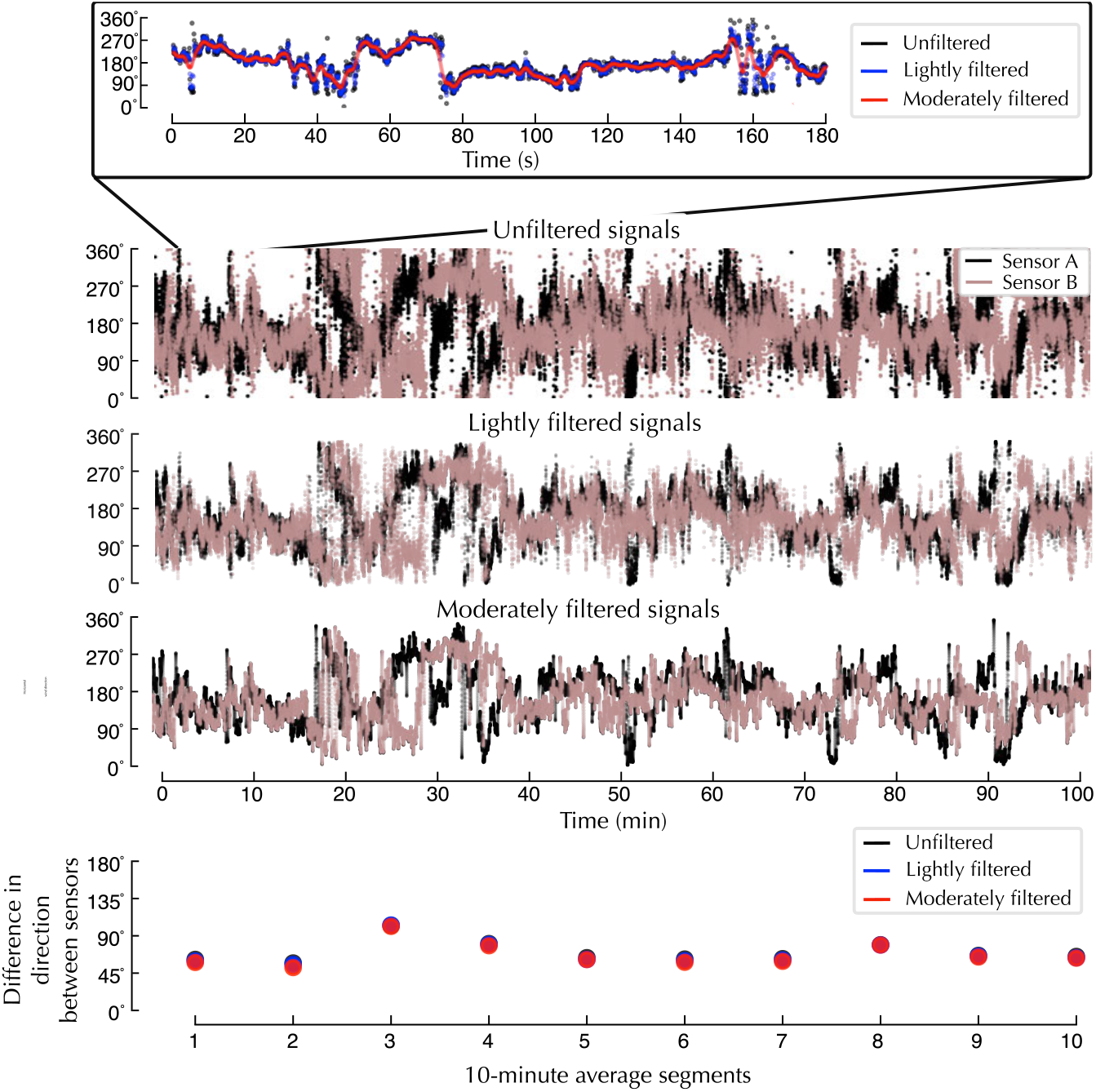
Signal smoothing does not have a strong effect on the mean differences between sensors. The bottom panel demonstrates the 10-minute mean difference between two sensors with unfiltered, lightly filtered and moderately filtered signals. In most cases, the computed differences were negligible, as can be seen by the overlapping points.

**Figure S5:**
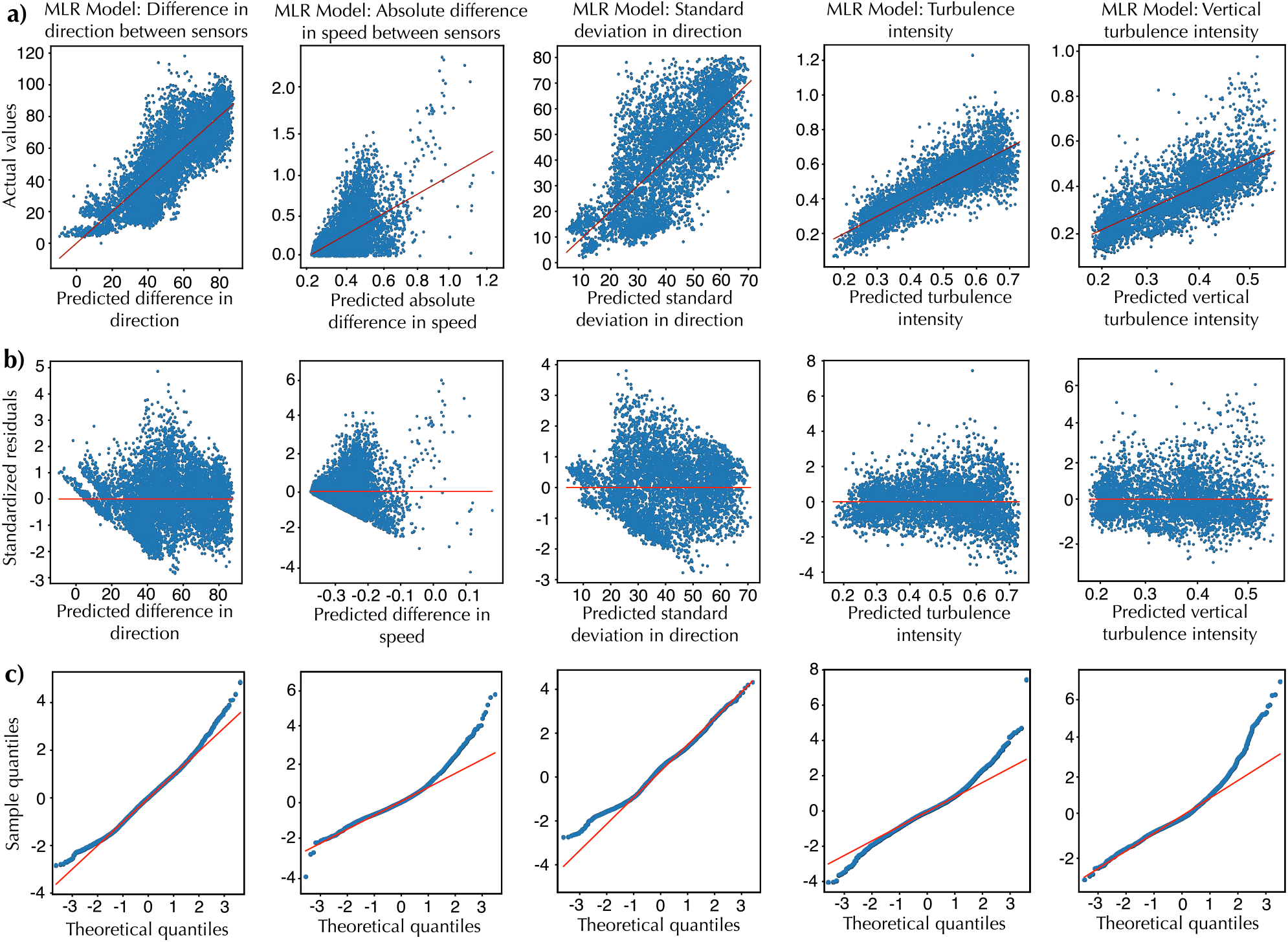
Diagnostic plots for the multiple linear regressions corresponding to the table values listed in Tables 1 & 2. A) actual values versus predicted values, B) standardized residuals versus predicted values, and C) Q-Q plot. Each column represents a MLR by the order it appears in the text.

**Figure S6:**
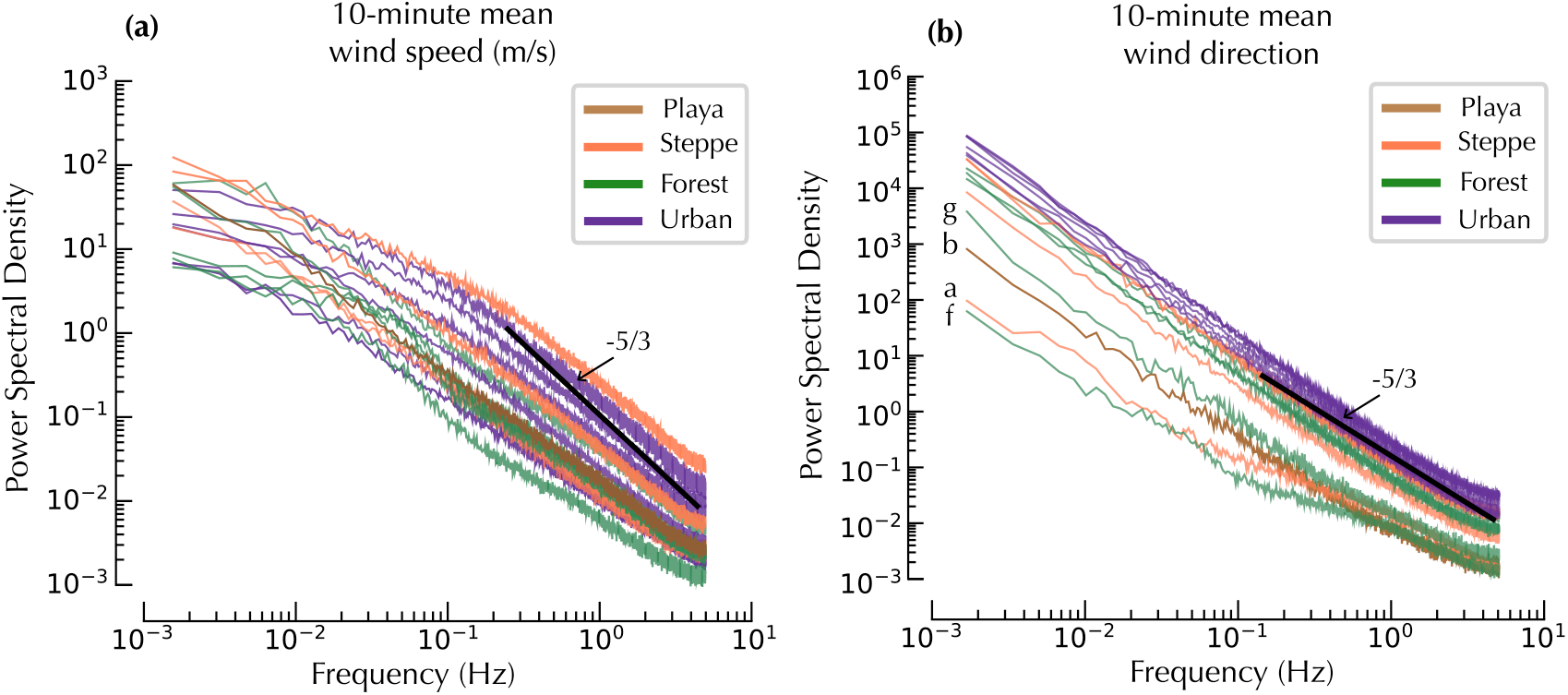
Frequency analysis: We assessed the variation of wind speed and direction with respect to the frequency domain by finding the mean power spectral density (PSD) of all 10-minute intervals across each time series. No significant differences were found at that time scale. A) 10-minute mean wind speed and B) 10-minute mean wind direction where a,b,c and d correspond to days with very little variation in wind direction (Supplemental Figures S1 & S2).

**Figure S7:**
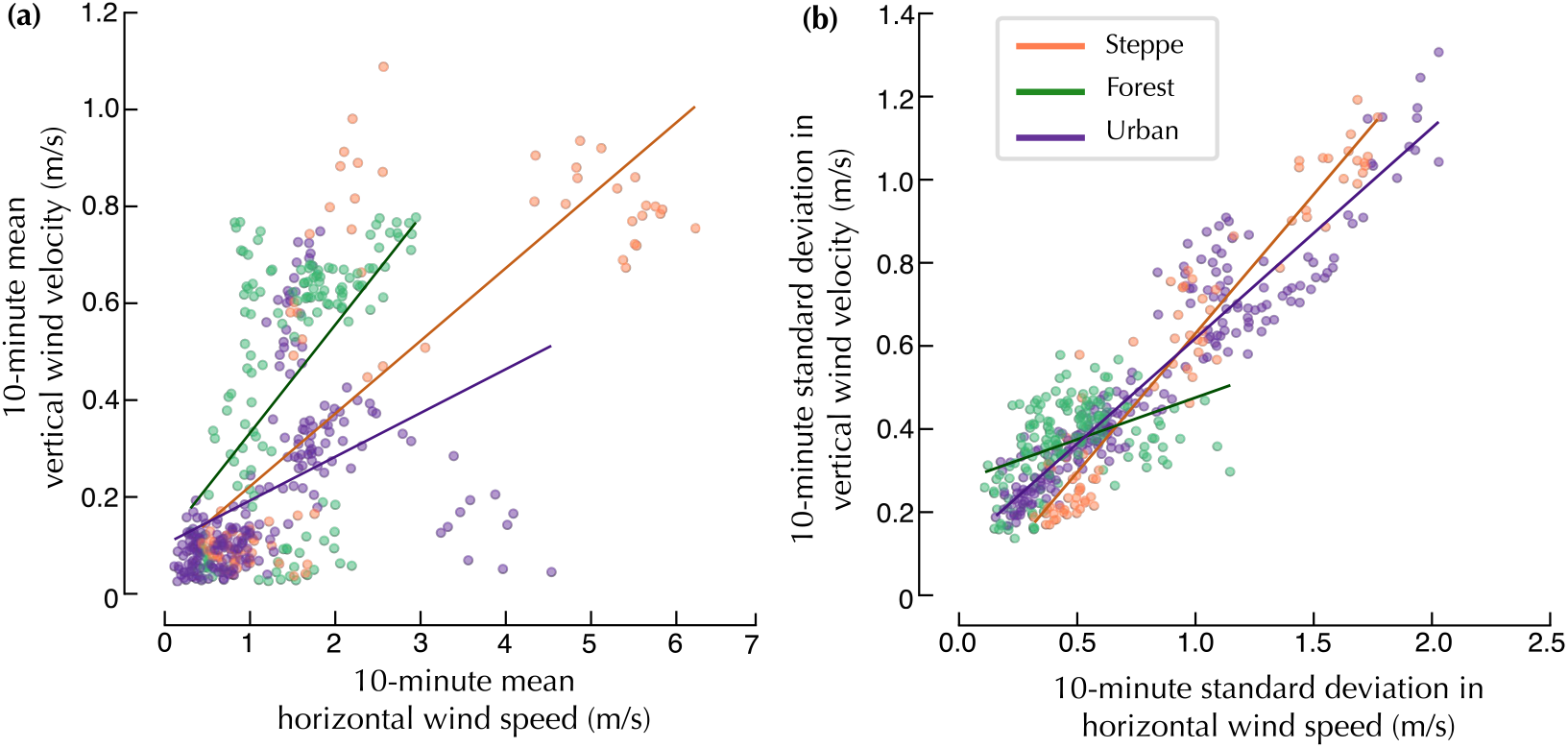
Mean vertical wind speed and *σ*_*w*_ generally increase with horizontal wind speeds. A) 10-Minute mean vertical velocity as a function of mean horizontal wind speed. B) 10-Minute *σ*_*w*_ as a function of *σ*_*h*_.

## Notes

### Competing Interest Statement

The authors have declared no competing interest.

